# Deoxyribonucleotide incorporation reshapes mRNA design beyond canonical ribonucleotide boundaries

**DOI:** 10.64898/2026.07.09.737403

**Authors:** Xiaoyan Ding, Riu Liao, Giovana Bavia Bampi, Dandan Zhang, Shan Guan, Joseph Rosenecker

## Abstract

Messenger RNA (mRNA) is canonically composed of ribonucleotides, with sporadic incorporation of deoxyribonucleotides into natural RNA transcripts being traditionally regarded as a rare, deleterious error arising from transcriptional infidelity. Here, we challenge this paradigm by demonstrating controlled partial substitution of ribonucleotides with deoxyribonucleotides during in vitro transcription (IVT) generates intact, stable and fully translationally competent IVT-mRNA. Unexpectedly, chimeric DNA-RNA backbone modification exhibits markedly enhanced IVT-mRNA translation several fold across multiple cell types and in vivo via diverse dosing routes relative to their ribonucleotide-based counterparts. 25% substitution of cytidine triphosphate with deoxycytidine triphosphate achieved best-performing translational output, surpassing the current gold-standard N1-methylpseudouridine (m1Ψ)-modified IVT-mRNA in a B16-OVA tumor vaccination model. These findings identify nucleotide class composition as a previously unrecognized parameter governing IVT-mRNA function and establish hybrid ribonucleotide-deoxyribonucleotide backbone engineering as a versatile strategy to expand the chemical space for next-generation mRNA therapeutics.

## Introduction

The clinical success of in vitro transcribed messenger RNA (IVT-mRNA) therapeutics, which has transformed modern medicine with unprecedented speed and flexibility(*1–4*), stems from the 2023 Nobel Prize-winning discovery that uridine-to-pseudouridine substitution markedly attenuates immunogenicity while boosting translation(*5*), revolutionizing IVT-mRNA from a fragile and immunostimulatory molecule into a clinically viable therapeutic platform(*6, 7*). Yet despite these advances, the chemical space explored for mRNA engineering remains remarkably constrained(*8*). Current optimization strategies focus predominantly on modifying the ribonucleotide building blocks—most notably through N^1^-methylpseudouridine (m1Ψ) incorporation(*9, 10*). Although revolutionary, these approaches remain grounded in the conventional ribonucleotide framework of mRNA, focusing on canonical RNA building block modifications while alternative nucleotide chemistries, such as DNA nucleotides incorporation, remain unexplored for IVT mRNA engineering.

The canonical distinction between RNA and DNA rests on the nature of their sugar backbone: ribonucleotides versus 2′-deoxyribonucleotides. This chemical difference underpins the fundamental separation of biological information storage and expression, with DNA serving as the stable genetic repository and RNA functioning as an intermediary molecule in gene regulation and transient protein synthesis(*11, 12*). Consequently, the presence of deoxyribonucleotides within RNA has generally been considered an aberration. Indeed, quantitative analyses have identified deoxyribonucleotides in cellular mRNAs, rRNAs, and tRNAs at frequencies ranging from approximately one deoxyribonucleotide per 10^3^–10^4^ ribonucleotides in organisms including *Escherichia coli*, *Saccharomyces cerevisiae*, mammalian cells, and RNA viruses(*13*). Accumulating evidences suggest that 2′-deoxyribonucleotides can be aberrantly incorporated into RNA. These sporadic incorporations have generally been attributed to polymerase infidelity or nucleotide pool imbalance and are therefore regarded as a rare and potentially deleterious event, presumed to compromise RNA structure, conformational flexibility, or translation rather than to confer any benefit(*14–17*).

This prevailing view, however, may have obscured a more nuanced reality. Dengue virus has been reported to contain comparatively elevated levels of deoxyribonucleotide incorporation within genomic RNA(*13*), suggesting the RNA molecules may tolerate a broader range of nucleotide compositions than traditionally assumed. Nevertheless, studies examining deoxyribonucleotides in RNA have primarily focused on naturally occurring mis-incorporation events and terminal modifications, particularly within poly(A) tails, leaving the biological and translational consequences of internal deoxyribonucleotide substitutions largely unexplored(*18, 19*).These pioneering studies suggest that the chemical composition of RNA backbone may serve as a functional determinant rather than merely a neutral structural feature. Unfortunately, whether systematic internal incorporation of deoxyribonucleotides throughout full-length IVT-mRNA is compatible with stable structure, or whether such modification can be harnessed to modulate protein expression, remains entirely unknown.

To this end, we investigated whether the in vitro transcription machinery could tolerate substitution of ribonucleotides with deoxyribonucleotides and how the resulting hybrid IVT-mRNA with altered physicochemical properties modulates translation profiles. We surprisingly found that deoxyribonucleotide can be incorporated during in vitro transcription and yielding intact, fully translation-competent IVT mRNA. More unexpectedly, certain substitution patterns, particularly 25% replacement of cytidine triphosphate (CTP) with deoxycytidine triphosphate (dCTP), consistently and significantly approached or even surpassed protein expression to levels that of the gold-standard m1Ψ[modified counterpart in both in vitro and in vivo models. The functional benefits of this hybrid backbone strategy extended beyond reporter assays. In a B16[OVA melanoma model, 25% dCTP[modified mRNA outperforms unmodified or m1Ψ[modified counterparts, positioning this strategy as a viable and potentially superior alternative to existing m1Ψ[modification approaches.

From a mechanistic standpoint, we speculate that the introduction of deoxyribonucleotides may alter RNA secondary structure, modulate ribosomal elongation kinetics, or affect interactions with RNA surveillance and decay pathways(*20*). The observation suggests a nuanced interplay between backbone chemistry and translational machinery, rather than a simple stability[based effect. Future structural and ribosome[profiling studies will be required to fully elucidate the underlying mechanisms.

Conceptually, our work expands the chemical space available for mRNA engineering beyond the ribose[based modifications that have dominated the field. By demonstrating that partial replacement of ribonucleotides with deoxyribonucleotides is not only tolerated but can be beneficial, we introduce nucleotide class composition as a previously unrecognized parameter governing mRNA function. This opens a new dimension for mRNA design, orthogonal to existing strategies such as codon optimization, UTR engineering, and nucleoside modification. It also raises broader questions about the evolutionary and biochemical boundaries of nucleic acid function, suggesting that the physicochemical constraints on primitive genetic systems may have been more flexible than previously appreciated.

## Results

### 1. Deoxynucleotide incorporation is feasible and generates intact IVT-mRNA transcripts

T7 RNA polymerase exhibits a strong preference for ribonucleoside triphosphates (NTPs) over deoxyribonucleoside triphosphates (dNTPs). However, the degree of substrate discrimination has been reported to vary among enzyme preparations from different commercial sources(*21, 22*). To determine whether the T7 RNA polymerase used in this study can incorporate deoxyribonucleotides into IVT-mRNA, we performed in vitro transcription using a linearized firefly luciferase (FLuc) template in which individual NTPs were progressively substituted with their corresponding dNTP counterparts.

Increasing the proportion of dNTPs resulted in a progressive reduction in IVT-mRNA yield (Supplementary Table 1), indicating that deoxyribonucleotide incorporation is associated with reduced transcriptional efficiency. Complete replacement of any one of the four ribonucleotides with its deoxy counterparts nearly abolished RNA synthesis entirely (Supplementary Table 1). Despite this reduction in yield, transcripts generated with 5%, 25%, or 50% dNTP substitution displayed comparable electrophoretic profiles, with no detectable evidence of degradation or truncation (Figure 1A). These findings indicate that, although extensive deoxynucleotide substitution impairs IVT-mRNA production, partial substitution remains compatible with the generation of intact, full-length transcripts.

**Figure 1.**
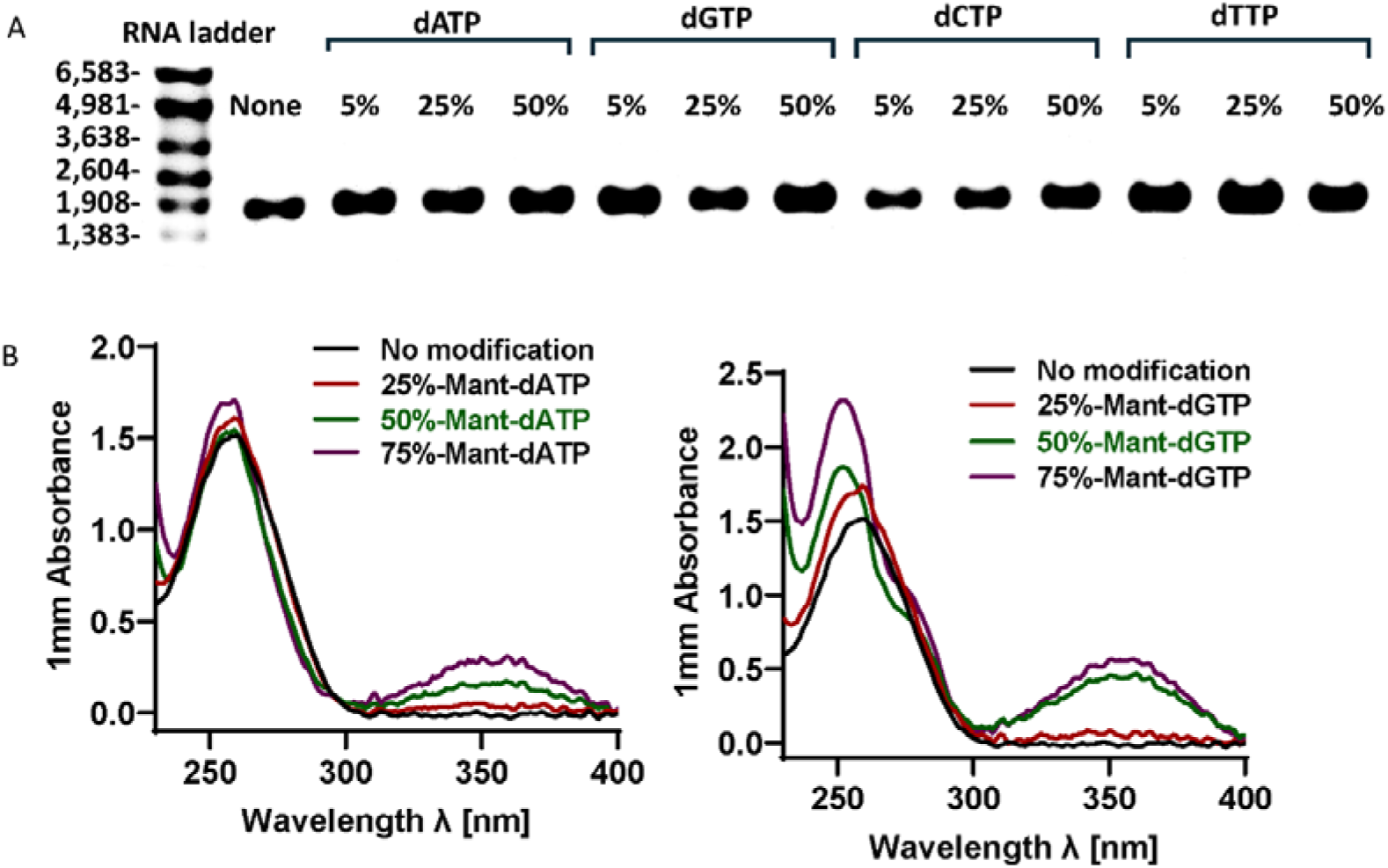
*In vitro* transcription of 2′-deoxyribonucleoside-incorporated mRNA by T7 polymerase. (A) Aliquots of in vitro–transcribed Firefly Luciferase (Fluc) containing no or 5%-, 25%-, 50%-dATP, dCTP, dGTP, or dTTP were separated in a denaturing 1% agarose gel. (B) Spectroscopic properties of 25%, 50%, 75%-Mant-dATP or 25%, 50%, 75%-Mant-dGTP-incoparated mRNA. The concentration of all RNA candidates is approximately 500 ng/µL, with each sample being repeated three times. The values shown in the figure represent the average of the three repetitions.

To directly verify deoxyribonucleotide incorporation, we employed two fluorescent dNTP analogs, 3′-O-(N-methylanthraniloyl)-2′-deoxyadenosine-5′-triphosphate (Mant-dATP) and 3′-O-(N-methylanthraniloyl)-2′-deoxyguanosine-5′-triphosphate (Mant-dGTP). Unlike canonical nucleotides, these analogs exhibit a characteristic absorbance peak at 350 nm (Supplementary Table 2). Following in vitro transcription and purification, transcripts synthesized in the presence of Mant-dATP or Mant-dGTP displayed a dose-dependent increase in absorbance at 350 nm, demonstrating successful incorporation of the fluorescent deoxyribonucleotide analogs into the IVT-mRNA products (Figure 1B).

Together, these results demonstrate that the T7 RNA polymerase used in this study tolerates partial deoxyribonucleotide substitution and it is possible to generate intact deoxy-modified mRNA transcripts, thereby establishing the experimental foundation for subsequent functional analyses.

### 2. dCTP incorporation enhances IVT-mRNA translation in cultured cells

To evaluate the functional consequences of partial deoxynucleotide incorporation, IVT-mRNA encoding firefly luciferase (Fluc) was synthesized with graded substitution of individual ribonucleotides (rNTPs) with the corresponding deoxyribonucleotides and assessed for translational activity in cultured cells.

An initial screening in DC2.4 cells identified several substitution conditions that enhanced luciferase expression relative to unmodified IVT-mRNA, with 25% 2′-deoxycytidine-5′-triphosphate (25% dCTP), 5% 2′-deoxythymidine-5′-triphosphate (5% dTTP), and 25% dTTP substitution yielding the most pronounced improvements (Figure 2A). To assess whether this effect was cell type–dependent, the top-performing candidates from the DC2.4 screen were further evaluated in A549 cells. Consistently, 25% dCTP-modified mRNA produced the highest luciferase expression (Figure 2B). A similar enhancement was also observed in HEK293T cells (Figure 2C), indicating that the effect is reproducible across both immune- and epithelial-derived cell lines.

**Figure 2.**
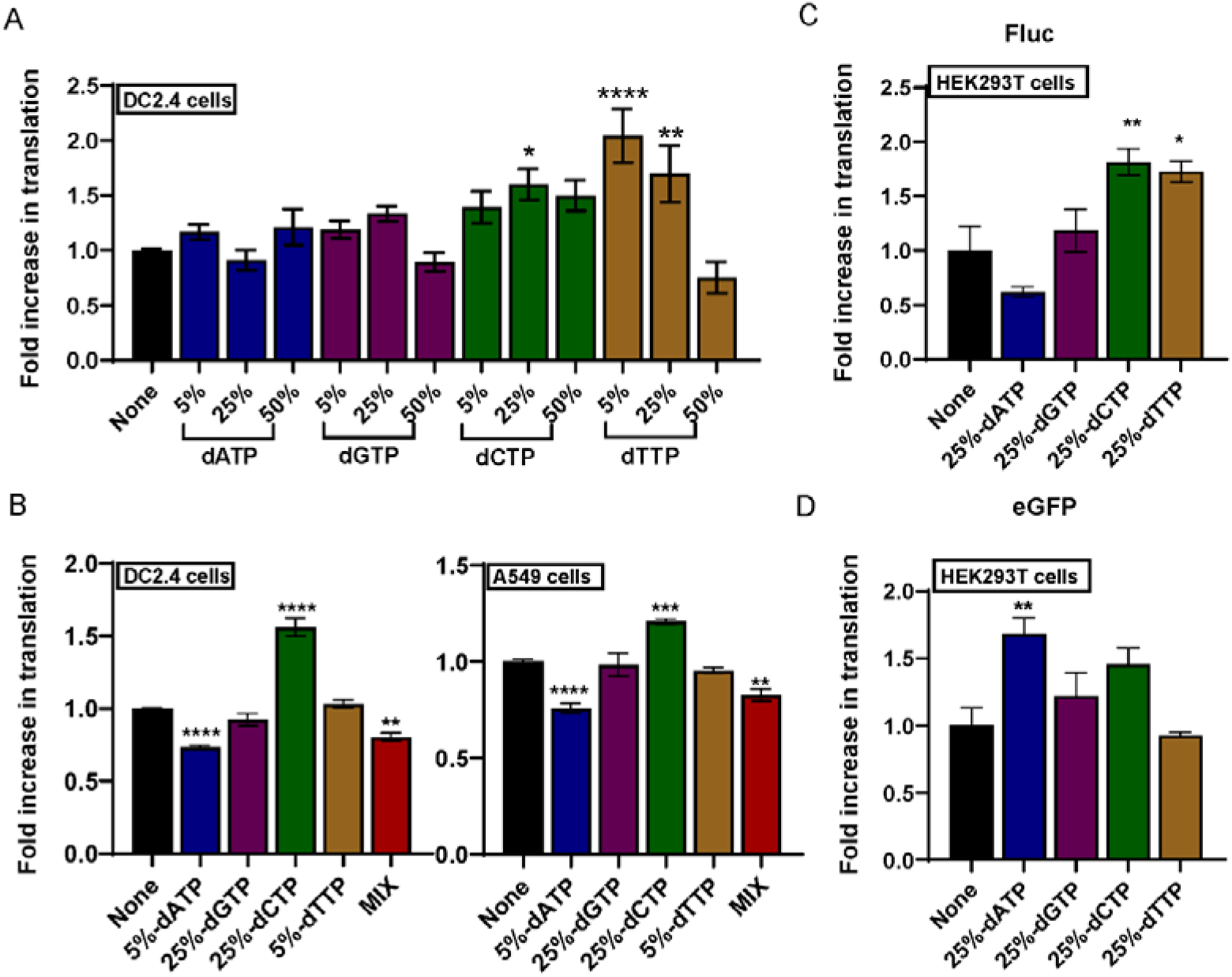
Cellular assessment of translation efficiency 24 h post-transfection of different *in-vitro* synthesized mRNA containing incorporated dNTP. (A) No or 5%, 25% and 50% dNTP incorporated into luciferase-expressing mRNA were transfected into DC2.4 cells. Firefly luciferase (Fluc) expression levels were measured 24 hours after transfection of 100 ng RNA encoding the Fluc reporter into DC2.4 cells using Lipofectamine™ RNAiMAX. Error bars indicate the standard error of the mean (SEM) from n = 15 replicates. (B) Top candidates identified in (A) were further evaluated in different cell types, including A549 cells and DC2.4 cells. mRNA was delivered using lipid nanoparticles (LNPs). Firefly luciferase (Fluc) expression levels measured 24 hours after transfection of 100 ng RNA encoding the Fluc reporter into DC2.4 cells or A549 cells. Error bars indicate the standard error of the mean (SEM) from n = 6 replicates. (C) Fluc expression after different 25%-dNTP-substituted mRNAs encoding Fluc were transfected into HEK293T cells. Error bars indicate the standard error of the mean (SEM) from n = 4 replicates. (D) eGFP expression after different 25%-dNTP-substituted mRNAs encoding Fluc were transfected into HEK293T cells. Error bars indicate the standard error of the mean (SEM) from n = 5 replicates. HEK293 T cells were seeded at about 2.5 × 10^4^ per well. Fluc and eGFP expression levels measured 24 hours after transfection of 200 ng mRNA encoding the Fluc reporter or eGFP reporter into HEK293T cells using Lipofectamine™3000. Luciferase activity was quantified using the Pierce™ Firefly Luciferase Glow Assay Kit. Luminescence intensity and fluorescence were measured with a FLUOstar Omega microplate reader (BMG Labtech). Approximately 4 × 10L DC 2.5 cells and 2.5× 10L A549 Cells were seeded per well. None: Unmodified RNA (no deoxynucleotide substitutions). “MIX” represents: 5%-dATP, 5%-dTTP, 25%-dCTP and 25%-dGTP substituted. Data are presented as mean ± SEM.

Finally, to determine whether this phenomenon extends beyond a single reporter system, enhanced Green Fluorescent Protein (eGFP) mRNA containing 25% dCTP substitution was tested and likewise exhibited increased protein expression compared to unmodified mRNA (Figure 2D), demonstrating that the translational enhancement is not restricted to a single coding sequence.

### 3. dCTP-modified IVT-mRNA enhances protein expression *in vivo*

We next examined whether the translational advantage of dCTP-modified IVT-mRNA was retained in vivo after intramuscular or inhaled administration. Following intramuscular delivery, 25% dCTP-modified IVT-mRNA generated stronger whole-body luciferase signals than WT IVT-mRNA at both 6 h and 24 h, while the 6 h signal remained lower than that induced by N^1^-methylpseudouridine (m1Ψ)-modified IVT-mRNA (Figure 3A-D). At 24 h, 25% dCTP produced luciferase expression comparable to or higher than m1Ψ, with a similar trend observed in the liver ex vivo, where 25% dCTP showed higher liver-associated bioluminescence than WT (no ribonucleotide substitution) and was comparable to m1Ψ (Figure 3E andF). After inhaled administration, 25% dCTP further increased pulmonary luciferase expression relative to WT at both 6 h and 24 h, and its whole-body signal was comparable to or higher than m1Ψ (Figure 3G-J). Ex vivo lung imaging confirmed that 25% dCTP markedly enhanced lung-localized expression compared with WT and reached a level similar to m1Ψ (Figure 3K and L). Histological analysis of major organs showed no apparent tissue injury after 25% dCTP vaccination, with tissue morphology comparable to WT and m1Ψ groups (Supplementary Figure 1). These results indicate that partial dCTP incorporation improves in vivo mRNA translation across distinct delivery routes, with expression levels approaching or exceeding those achieved by the widely used m1Ψ modification in route- and tissue-dependent contexts.

**Figure 3.**
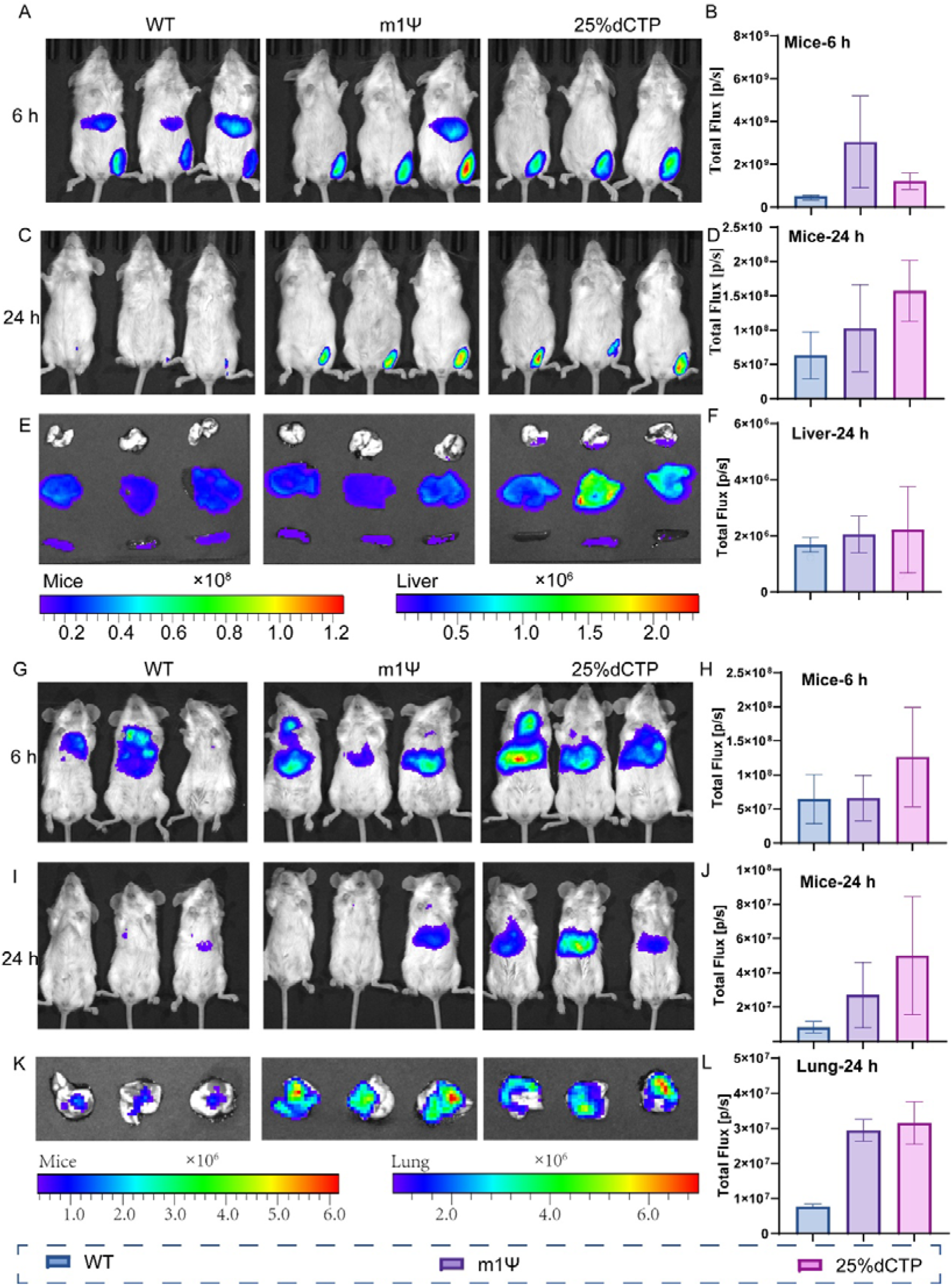
In vivo and organ biodistribution profiles following different administration routes. (A-D) In vivo bioluminescence imaging (BLI) of mice receiving WT, m1Ψ, or 25%dCTP loaded with reporter mRNA. Representative whole-body images at 6 h (A) and 24 h (C) post-injection, with corresponding total flux quantification (B, D). (E, F) Ex vivo organ imaging of liver harvested at 24 h post-administration. Representative radiant efficiency heatmaps (E) and quantitative total flux (F) demonstrate detectable reporter expression across all groups. (G-J) In vivo BLI of lung-region signals at 6 h (G) and 24 h (I). Quantification (H, J) reveals markedly increased pulmonary luminescence in 25%dCTP-LNP–treated animals relative to WT and m1Ψ LNPs, particularly at 24 h. (K, L) Ex vivo lung imaging at 24 h. Representative images (K) and total flux analysis (L). Data are presented as mean ± SD. (n = 3 per group).

### 4. dCTP-modified IVT-mRNA improves antitumor immunity and therapeutic efficacy

To test whether enhanced translation improved vaccine efficacy, we evaluated 25% dCTP-modified IVT-mRNA in a prophylactic B16-OVA tumor challenge model(*23, 24*) (Figure 4A). Body-weight monitoring showed no additional systemic burden in the 25% dCTP group compared with the other mRNA-treated groups during the observation period, whereas mice treated by PBS control an exhibited significant weight loss since Day 38 (Figure 4B). Compared with WT and m1Ψ-modified IVT-mRNA, 25% dCTP induced higher OVA-specific IgG titers after immunization (Figure 4C). Flow-cytometric analysis further revealed that 25% dCTP increased the frequencies of CD44□CD62L□CD8□ T cells, Granzyme B□ CD8□ T cells and IFN-γ□ CD8□ T subsets relative to WT and m1Ψ counterparts (Figure 4D-F and Supplementary Figure 2A-C). As vividly depicted in Figure 4G, tumor sections from the 25% dCTP group showed stronger CD3□, CD8□ and CD11c□ immune-cell infiltration than those from WT- and m1Ψ-treated counterparts. These immune changes were associated with reduced endpoint tumor burden and delayed tumor growth in the 25% dCTP group compared to PBS, WT and m1Ψ groups (Figure 4I and 4J). Survival analysis showed a trend towards improved outcome after 25% dCTP vaccination, although the difference among mRNA-treated groups was not significant (Figure 4H).

**Figure 4.**
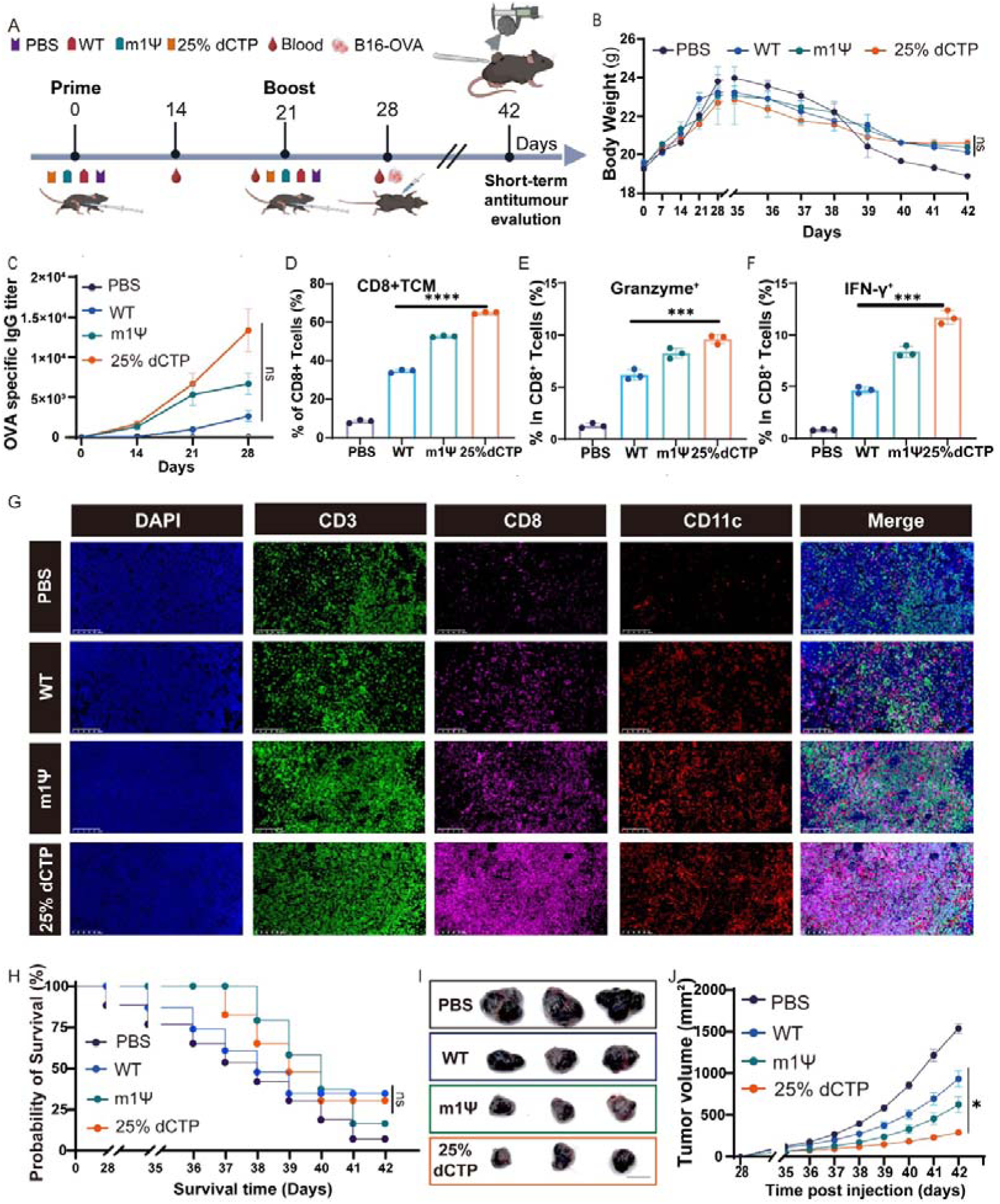
25% dCTP immunization induces potent anti-tumor immunity and improves survival. (A) Schematic illustration of the vaccination and challenge schedule. Mice were immunized on days 0 and 21 and challenged with B16-OVA cells on day 28, followed by short-term antitumor evaluation on day 42. Blood was collected periodically. (B) Body weight changes of mice in different treatment groups (PBS, WT, m1Ψ, 25% dCTP) over the course of the experiment (n=10 biologically independent animals). (C) Serum OVA-specific IgG titers were measured at the indicated time points after immunization. (D) The proportion of central memory T cells (TCM, CD44+CD62L+) in CD8+ T cells in inguinal lymph nodes of mice vaccinated by indicated formulations 28 days post-prime. (E and F) Quantification of granzyme B□ CD8□ T cells (CD8LGranzyme B□) (E) and IFN-γ□ CD8L T cells (CD8LIFN-γ□) (F) in lymphocytes isolated from inguinal lymph nodes at 28 days post-prime. (G) Immunofluorescence staining of inguinal lymph-node sections showing DAPI, CD3, CD8, and CD11c signals and merged images. CD3 (T cells, green), CD8 (cytotoxic T, pink) and CD11c (dendritic cells, red). Nuclei are counterstained with DAPI (blue). Scale bar, 100 µm. (H) Survival curves of mice following the various treatments. (I) Representative images of excised B16-OVA melanoma tumors collected at endpoint. (J) Tumor growth curves following different treatments. Data are presented as mean ± SD.

Histological analyses further supported the antitumor effect of 25% dCTP-modified IVT-mRNA. Ki67 immunohistochemistry showed markedly lower tumor-cell proliferation in the 25% dCTP group than in PBS, WT and m1Ψ groups (Supplementary Figure 2D and 2E). H&E staining also showed reduced tumor thickness after 25% dCTP vaccination, consistent with attenuated tumor progression at the tissue level (Supplementary Figure 2F and 2G). Together, these data indicate that partial dCTP incorporation enhances the humoral and CD8□ T-cell responses elicited by IVT-mRNA vaccination and improves short-term antitumor protection relative to WT and m1Ψ-modified controls.

## Discussion

In this study, we demonstrate that controlled partial substitution of ribonucleotides with deoxyribonucleotides during in vitro transcription (IVT) can generate full-length IVT-mRNAs that remain intact and translationally competent. Although T7 RNA polymerase strongly favours canonical NTPs, efficient deoxyribonucleotide incorporation was achieved by titrating the deoxy-substrate into the IVT reaction rather than replacing the corresponding ribonucleotide completely. Specifically, a defined fraction of CTP was replaced with dCTP while maintaining the remaining NTPs and the total cytidine nucleotide input, allowing T7 RNA polymerase to incorporate dCTP without substantially compromising transcript elongation. Among the tested substitution levels, 25% dCTP provided the best balance between full-length transcript production, mRNA stability and translational output. These findings extend the functional limits of the IVT-based mRNA synthesis and provide an experimental basis for exploring IVT-mRNA engineering beyond the classic ribonucleotide paradigm.

Current IVT-mRNA engineering strategies, most notably the incorporation of N¹-methylpseudouridine, operate largely within a fixed ribonucleotide backbone, representing optimization within a chemically constrained space(*25, 26*). By contrast, the introduction of deoxyribonucleotides as compositional variables expands the chemical space of IVT-mRNA beyond ribose-based modifications. This shift from intra-backbone modification to backbone-class heterogeneity suggests that nucleotide composition may constitute an additional, previously underappreciated determinant of mRNA translational output, extending current design principles for synthetic IVT-mRNA.

Across multiple reporter systems (Fluc and eGFP) and experimental conditions, we observed consistent enhancement of translational output, indicating that the effect is not restricted to a specific coding sequence context. In particular, partial substitution with 25% dCTP resulted in reproducible increases in protein expression across cell types, suggesting that nucleotide composition can influence mRNA performance in a substitution-level-dependent manner. This strategy maintained efficient in vivo translation across different delivery routes and was associated with improved performance in a tumor vaccination model. These observations suggest that alterations in nucleotide backbone composition can improve RNA translation and may provide a tractable strategy for optimizing mRNA-based therapeutics.

The molecular mechanisms underlying these effects remain to be fully elucidated. Plausible contributing factors include changes in RNA secondary structure and conformational dynamics, modulation of ribosomal elongation kinetics, and alterations in transcript stability. In addition, RNA surveillance and degradation pathways, including RNase-mediated processes, may contribute to the observed differences. Recent work by Bérouti et al. showed that pseudouridine-containing RNA avoids innate immune detection by impairing endolysosomal RNA processing and limiting generation of TLR7/8-stimulatory RNA fragments(*27*). This mechanism provides an important reference point for interpreting modified mRNA activity. Notably, a very recent landmark study has systematically characterized N4-acetylcytidine, an untested synthetic mRNA modification that confers higher mRNA translation yield and fidelity than the widely used m1Ψ modification by moderately slowing elongation to avoid excessive ribosome collisions, downstream ribosome-associated quality control activation and +1 frameshifting while retaining equivalent RNase T2 resistance to suppress innate immune activation(*28*).

We hypothesize that 25% dCTP-modified IVT mRNA may represent an optimization strategy distinct from that of m1Ψ-modified mRNA. Although its protein expression was comparable to or even higher than that of m1Ψ-modified mRNA, 25% dCTP modification probably did not simply mimic the immune-silent phenotype associated with Ψ/m1Ψ modifications. Instead, in the B16-OVA vaccination model, 25% dCTP-modified mRNA retained immunostimulatory activity and induced stronger antigen-specific T-cell responses, suggesting that enhanced protein expression was achieved without compromising adaptive immune activation. The increased expression may involve mechanisms related to improved mRNA stability and ribosome loading efficiency, although further studies are required to define the underlying molecular basis. Thus, partial dCTP incorporation may complement Ψ/m1Ψ-based strategies by improving translational performance without fully suppressing vaccine-relevant immune activation.

Finally, the observed tolerance of transcript and translation machinery toward partial deoxyribonucleotide incorporation raises a broader conceptual point regarding nucleic acid evolution. Rather than implying specific adaptive advantages, this observation is consistent with a scenario in which early nucleic acid systems were not strictly constrained to a uniform ribose-based backbone, but may have tolerated mixed or partially heterogeneous nucleotide chemistries while maintaining functional output. Such biochemical permissiveness could, in principle, have expanded the sequence–structure–function space accessible to primitive replicating systems, prior to the emergence of highly optimized substrate-selective polymerases. While speculative, this perspective is consistent with the notion that the physicochemical constraints governing nucleic acid function may have been more flexible in primitive molecular environments, and may still be partially reflected in the residual substrate tolerance observed in contemporary enzymatic systems.

## Conclusion

In summary, this study identifies deoxyribonucleotide incorporation as a previously unrecognized compositional parameter influencing mRNA function, inviting a re□evaluation of the compositional plasticity of mRNA. Our findings establish hybrid ribonucleotide□deoxyribonucleotide backbone engineering as a versatile strategy for enhancing IVT-mRNA translation that extends beyond conventional ribonucleotide-based modification paradigms, expanding the chemical space available for IVT-mRNA design and highlighting nucleotide class composition as an additional determinant of gene expression output. The convergence of deoxyribonucleotide chemistry with ribonucleotide macromolecules is expected to open new frontiers in the chemical optimization of mRNA therapeutics with tunable translational properties.

## Materials and Methods

### RNA design and synthesis

The DNA templates (sequences shown in Supplementary Table 3) comprised codon-optimized firefly luciferase (Fluc) or enhanced green fluorescent protein (eGFP) open reading frames (ORFs) flanked by previously optimized untranslated regions(*29, 30*). An A30L70 segmented poly(A) cassette (30A–GCATATGACT–70A, as described in WO2016005324A1) was incorporated at the 3′ end. All sequences were placed under the control of a T7 promoter (TAATACGACTCACTATA) and cloned into the pUC57 vector (GenScript), where they were synthesized and sequence-verified.

Before in vitro transcription (IVT), 1 µg of plasmid DNA was linearized using BsaI-HFv2 restriction enzyme (New England Biolabs, R3733S) for 1 h at 37°C, followed by purification using a QIAquick Gel Extraction Kit (Qiagen, 28706).

IVT-mRNA synthesis was performed as previously described(*29, 31*), using the HiScribe™ T7 High Yield RNA Synthesis Kit (New England Biolabs, E2040S) according to the manufacturer’s instructions with minor modifications. Briefly, 20 µL IVT reactions were assembled using linearized DNA templates containing a T7 promoter. For deoxynucleotide substitution experiments, ribonucleoside triphosphates (NTPs) were partially replaced with corresponding deoxyribonucleoside triphosphates (dNTPs: dATP, dCTP, dGTP, or dTTP) at defined ratios.

Co-transcriptional capping was achieved by addition of CleanCap® AG (TriLink Biotechnologies, N-7113) at a final concentration of 6 mM. Reactions were incubated at 37°C for 3 h, followed by DNase I treatment (Thermo Scientific™, EN0525) for 15 min. RNA was purified by lithium chloride (LiCl) precipitation. RNA concentration was measured using a NanoDrop spectrophotometer, and RNA integrity was assessed by 1% agarose gel electrophoresis.

Different percentages of mant-dATP and mant-dGTP substitutions were applied during FLUC mRNA synthesis following the same IVT procedure described above. In these experiments, only the indicated nucleoside triphosphates were replaced, while all other reaction components remained unchanged. For example, for 25% mant-dATP substitution, the reaction contained ATP and mant-dATP (Jena Bioscience, NU-203S) at final concentrations of 5.625 mM and 1.875 mM, respectively, while maintaining the total adenine nucleotide concentration constant.

Following transcription, RNA samples (500 ng/µL in nuclease-free water) were subjected to UV–Vis absorption analysis using a NanoDrop™ 2000/2000c spectrophotometer (Thermo Fisher Scientific, ND-2000C). Baseline correction was performed using the corresponding solvent before measurements.

### Cell culture, transfection and analysis

DC2.4 (Sigma-Aldrich, SCC142) were cultured in DMEM (ATCC, 30-2002) with 10% FBS (ATCC, 30-2020). When confluence reaches 90% the cells are split and seeded in a 96-well plate at 4×10^4 per well. When the cell confluence reaches 70%, then using Lipofectamine RNAiMAX Transfection Reagent (Invitrogen, Cat. No. 13778075), 100 ng mRNA per well is transfected. After 24 hours of co-incubation, remove the supernatant of the cells. And following the instructions of the Pierce™ Firefly Luciferase Glow Assay Kit (Thermo Fisher, 16176) to quantify the expression of firefly luciferase.

For HEK 293T cells culture, HEK 293T cells (ATCC, CRL-3216™) were cultured in DMEM (ATCC, 30-2002) with 10% FBS (ATCC, 30-2020). When confluence reaches 90% the cells are split and seeded in a 96-well plate at 2.5×10^4^ per well in 100 uL DMEM with 2% FBS. When the cell confluence reaches in 70%, using Lipofectamine™ 3000 Transfection Reagent (Invitrogen, Cat. No. L3000008) transfected 200 ng Fluc or eGFP mRNA per well. After 24 hours, following the instructions of the Firefly Luciferase HTS assay (MilliporeSigma, Cat.NO.SCT150) to quantify the expression of firefly luciferase.

For A549 cells culture, cells were cultured in DMEM (ATCC, 30-2002) with 8% FBS (ATCC, 30-2020). When confluence reaches 90% the cells were split and seed in a 96 well plate at 2.5×10^4^ per well in 100 uL DMEM with 2% FBS. When the cell confluence reaches in 70%, transfected 100 ng 100 ng LNP-RNA encoding the Fluc reporter into cells per well. After 24 hours, following the instruction of Firefly Luciferase HTS assay (MilliporeSigma, Cat.NO.SCT150) to quantify the expression of firefly luciferase.

Luminescence intensity and fluorescence were measured with a FLUOstar Omega microplate reader (BMG Labtech).

### Lipid nanoparticle preparation

SM-102 (MCE, 251104), DSPC (MCE, 261435), cholesterol (Sigma-Aldrich, 57-88-5), and DMG-PEG2000 (Avanti, 880151p-1g-A-025) were dissolved in ethanol at a molar ratio of 50:10:38.5:1.5. mRNA was diluted in 50 mM citrate buffer (pH 4.0) to a final concentration of 0.17 mg/mL. Lipid and mRNA solutions were mixed at a nitrogen-to-phosphate (N/P) ratio of 6:1 using the NanoAssemblr Ignite microfluidic system (Precision Nanosystems), with a total flow rate of 12 mL/min and an aqueous-to-organic phase flow rate ratio of 3:1. The resulting lipid nanoparticle (LNP)-encapsulated mRNA formulations were dialyzed against phosphate-buffered saline (PBS, pH 7.4) for 24 hours using dialysis tubing (Viskase, USA), and stored at 4°C until further use. Encapsulation efficiency was assessed using the Quant-iT RiboGreen RNA Assay Kit (Invitrogen, USA), and fluorescence was measured with a FLUOstar Omega microplate reader (BMG Labtech).

### Physicochemical characterization and IVT-mRNA encapsulation efficiency (EE) analysis

The particle size and zeta potential of Lipid nanoparticle–mRNA formulations were determined using a Zetasizer Nano ZS (Malvern Instruments) at 25 °C as previously described(*32*), while their morphology was examined by transmission electron microscopy (TEM, JEM-1400plus, JEOL Ltd., Japan). For TEM imaging, samples were negatively stained with 2% uranyl acetate. The viscosity of the formulations was measured using a HAAKETM MARSTM viscometer (Thermo Fisher Scientific, USA). To assess IVT-mRNA encapsulation efficiency (EE), Lipid nanoparticle–mRNA samples were dispersed in 1×Tris-EDTA (TE) buffer or lysed with 2% Triton X-100 and incubated at 37 °C for 30 min. RNA standard solutions were prepared at concentrations of 0 µg/mL, 0.02 µg/mL, 0.1 µg/mL, 0.5 µg/mL, and 1 µg/mL using 1×TE buffer. For fluorescence measurement, 100 µL of each sample or standard was mixed with 100 µL of RiboGreen reagent (1:200 dilution) in a clear-bottom black plate, incubated at room temperature for 3 min, and analyzed using a VarioskanTM LUX microplate reader (Thermo Fisher Scientific) at excitation and emission wavelengths of 480 nm and 520 nm.

### Animal studies

Ethical Compliance and Experimental Design: The animal research protocol was conducted in compliance with international laboratory animal care standards (Approval No. AMUWEC20230184) and authorized by the Institutional Animal Ethics Committee of TMMU. The experimental design incorporated five representative rodent models: (1) female Balb/c mice (6-8 weeks); (2) female C57BL/6 mice (6-8 weeks).

All subjects were housed in individually ventilated cages (IVC systems) under strict specific pathogen-free (SPF) conditions, maintaining controlled environment parameters: temperature 22 ± 1 °C, humidity 55 ± 5%, with 12 h circadian rhythm regulation. Sterilized feed 981 (autoclaved pellet diet) and acidified water (pH 2.5-3.0) were provided ad libitum. Following a 7-day acclimatization period involving daily health monitoring (body weight, coat condition, activity levels), animals underwent stratified randomization based on weight-matching principles before experimental interventions.

### B16-OVA Tumor Models

For the B16-OVA prophylactic tumour model, female C57BL/6 mice were randomly divided into four groups: PBS, WT IVT-mRNA, N1-methylpseudouridine-modified IVT-mRNA (m1Ψ) and 25% dCTP-modified IVT-mRNA. The mRNAs encoded the OVA antigen (Suppmentary Table 3), and the dose of OVA mRNA was 3 μg per mouse. Mice were immunized on day 0 and boosted on day 21 with the corresponding formulations. PBS-treated mice received an equal volume of PBS as control. Blood samples were collected on days 14, 21 and 28 for serum OVA-specific IgG analysis. On day 28, mice were challenged with 1.0 × 10□B16-OVA cells by subcutaneous injection into the right dorsal flank. Body weight and tumour growth were monitored after tumour inoculation, and short-term antitumour efficacy was evaluated on day 42. Tumour length and width were measured using a digital caliper, and tumour volume was calculated using the formula: tumour volume = L × S²/2, where L represents the longest diameter and S represents the shortest diameter. Mice were euthanized when they reached the predefined humane endpoint, and tumours and relevant tissues were collected for further analysis.

For immune profiling, inguinal lymph nodes were collected from parallel cohorts at day 28 after prime immunization. Single-cell suspensions were prepared and analysed by flow cytometry to determine the frequencies of CD44□CD62L□CD8□T cells, Granzyme B□CD8□T cells and IFN-γ□ CD8□T cells. At the endpoint, excised tumours were imaged and processed for immunofluorescence staining, Ki67 immunohistochemistry and H&E staining to evaluate immune-cell infiltration, tumour-cell proliferation and histopathological tumour progression.

### Histopathology analysis

24 h post-administration of intramuscular injection, major organs of mice, including lungs, livers, spleens and kidneys, were collected and fixed overnight in 4 % paraformaldehyde at RT. The fixed organs were dehydrated, embedded in paraffin and sliced into 5 μm thickness. The lung sections were stained with hematoxylin and eosin (H&E) and visualized using a light microscope (Eclipse Ci-L, Nikon).

### Flow cytometry analysis

To detect the effector memory T Cells (TEM), central memory T cells (TCM) and tissue-resident memory T cells (TRM), pulmonary lymphocytes or splenocytes were prepared from immunized mice on 28 days post-immunization as previously described. Subsequently, cell samples were stained with the following antibodies in cell staining buffer for 40 min at 4 °C: anti-mouse-CD45-Brilliant Violet 421™ (Biolegend, 102133, 1:100), anti-mouse-CD3-Brilliant Violet 510™ (Biolegend, 100233, 1:100), anti-mouse-CD8-PE-Cy7 (Biolegend, 100721, 1:100), anti-mouse-CD62L-PE (553151, BD Biosciences, 1:100), anti-mouse-CD44-APC (559250, BD Biosciences, 1:100), anti-mouse-IFN-γ-BV 650 (48-7311-82, Invitrogen, 1:50), anti-mouse-Granzyme B-APC (Biolegend, 108723, 1:60).

### Histology, immunohistochemistry and immunofluorescence staining

Tumor tissues and major organs, including lung, kidney, liver and spleen, were collected at the indicated time points, fixed in 4% paraformaldehyde, embedded in paraffin and sectioned into 5-µm-thick slices. For histopathological analysis, sections were subjected to haematoxylin and eosin staining according to standard protocols and visualized using a light microscope (Eclipse Ci-L, Nikon).

For immunohistochemical analysis of tumor cell proliferation, paraffin-embedded tumor sections were deparaffinized, rehydrated and subjected to antigen retrieval. Endogenous peroxidase activity was blocked with 3% hydrogen peroxide, followed by blocking with serum or bovine serum albumin. Sections were then incubated with anti-Ki67 primary antibody overnight at 4 °C, followed by incubation with an HRP-conjugated secondary antibody. The chromogenic reaction was developed using a DAB Horseradish Peroxidase Color Development Kit (ZLI-9019, Beijing Zhongshan Golden Bridge Biotechnology Co., Ltd), and nuclei were counterstained with haematoxylin. Ki67 staining was imaged using a microscope and quantified by calculating the Ki67-positive area or positive-cell fraction in randomly selected fields.

For immunofluorescence staining, tumor sections were deparaffinized, rehydrated, subjected to antigen retrieval and blocked as described above. Sections were incubated with primary antibodies against CD3, CD8 and CD11c overnight at 4 °C, followed by incubation with species-matched fluorophore-conjugated secondary antibodies. Nuclei were counterstained with DAPI. Fluorescence images were acquired using a fluorescence microscope or confocal microscope, and the infiltration of CD3□, CD8□and CD11c□cells was quantified from representative tumor regions using ImageJ.

### Statistical Analysis

Statistical analysis was performed using Prism 9.0.0 (GraphPad Software). Dual comparisons were made using Welch’s t-test, and comparisons between multiple conditions were analyzed using analysis of variance (ANOVA) followed by the appropriate post-hoc tests. All of the tests were two-tailed. Differences were considered statistically significant when *P*<0.05.

## Author contributions

Jo.R., S.G., and X.D. coordinated the project. R.L., X.D., and G.B.B. conceived and designed the experiments. R.L., X.D., D.Z., and G.B.B. performed the experiments. X.D., R.L., and G.B.B. acquired, analyzed, and interpreted the data. Jo.R., S.G., X.D., R.L., and G.B.B. wrote the manuscript. Jo.R. and S.G. acquired funding. All authors read, reviewed, and approved the final manuscript.

## Data availability

The authors declare that the data supporting the findings of this study are available within this article and its supplementary information. Source data are provided with this manuscript.

## Ethics approval

All in vivo studies were approved by the Laboratory Animal Welfare and Ethics Committee of Third Military Medical University (AMUWEC20230184). They were performed in accordance with the institutional and national policies and guidelines for the use of laboratory animals.

## Supplementary Information

**Supplementary Table 1.** The concentration and purity of different percentages of dNTP-incorporated IVT-mRNA.

| Samples | Concentration<br>[ng/μL] | OD260/OD280 | OD260/OD230 |
| --- | --- | --- | --- |
| No Modification | 2443 | 2.15 | 2.16 |
| 25%-dATP | 1539 | 2.22 | 2.46 |
| 50%-dATP | 693 | 2.07 | 2.48 |
| 75%-dATP | 278 | 2.11 | 2.4 |
| 100%-dATP | 17.5 | 2.5 | 2.2 |
| 25%-dCTP | 1066 | 2.2 | 2.41 |
| 50%-dCTP | 1025 | 2.21 | 2.37 |
| 75%-dCTP | 767 | 2.22 | 2.35 |
| 100%-dCTP | 80.3 | 1.79 | 2.11 |
| 25%-dGTP | 1124 | 2.26 | 2.46 |
| 50%-dGTP | 588 | 2.15 | 2.37 |
| 75%-dGTP | 435 | 2.16 | 2.29 |
| 100%-dGTP | 109 | 1.76 | 2.38 |
| 25%-dTTP | 1128 | 2.17 | 2.38 |
| 50%-dTTP | 1174 | 2.22 | 2.42 |
| 75%-dTTP | 472 | 2.08 | 2.32 |
| 100%-dTTP | 112.4 | 1.74 | 2.22 |

**Supplementary Table 2.** Spectroscopic Properties of Different Nucleotides.

| Nucleotides | Spectroscopic Properties |
| --- | --- |
| ATP | $\lambda_{\text{max}}$ 259 nm, $\epsilon$ 15.1 L mmol<sup>-1</sup> cm<sup>-1</sup> (Tris-HCl pH 7.0) |
| GTP | $\lambda_{\text{max}}$ 252 nm, $\epsilon$ 14.2 L mmol<sup>-1</sup> cm<sup>-1</sup> (Tris-HCl pH 7.0) |
| UTP | $\lambda_{\text{max}}$ 262 nm, $\epsilon$ 9.8 L mmol<sup>-1</sup> cm<sup>-1</sup> (Tris-HCl pH 7.0) |
| CTP | $\lambda_{\text{max}}$ 271 nm, $\epsilon$ 8.9 L mmol<sup>-1</sup> cm<sup>-1</sup> (Tris-HCl pH 7.0) |
| Mant-dATP | $\lambda_{\text{max}}$ 255/355 nm, $\epsilon$ 23.3/5.8 L mmol<sup>-1</sup> cm<sup>-1</sup> (Tris-HCl pH 7.5), $\lambda_{\text{exc}}$ 355 nm, $\lambda_{\text{em}}$ 448 nm |
| Mant-dGTP | $\lambda_{\text{max}}$ 252/355 nm, $\epsilon$ 22.6/5.7 L mmol<sup>-1</sup> cm<sup>-1</sup> (Tris-HCl pH 7.5), $\lambda_{\text{exc}}$ 355 nm, $\lambda_{\text{em}}$ 448 nm |

**Supplementary Table 3.**
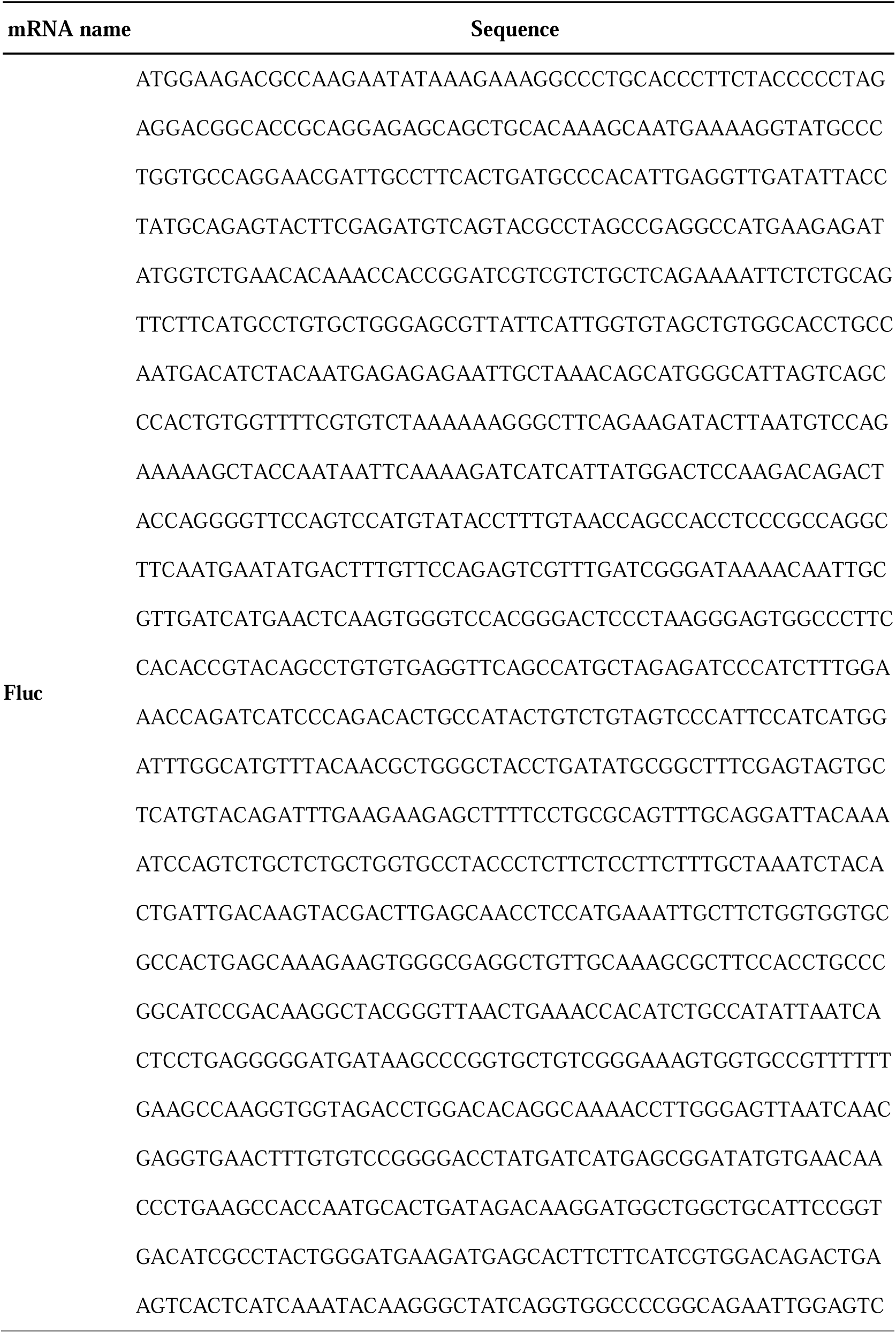
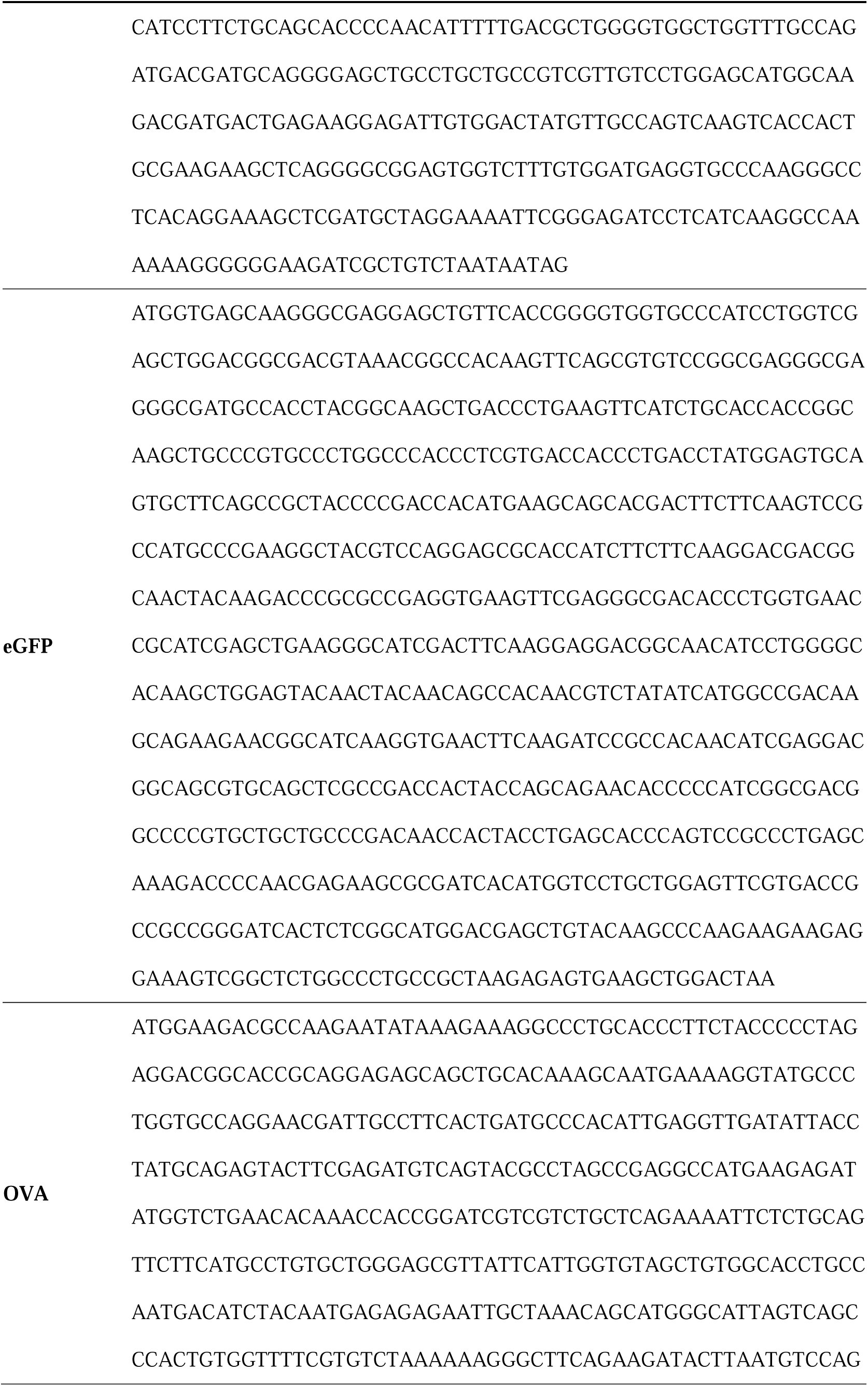
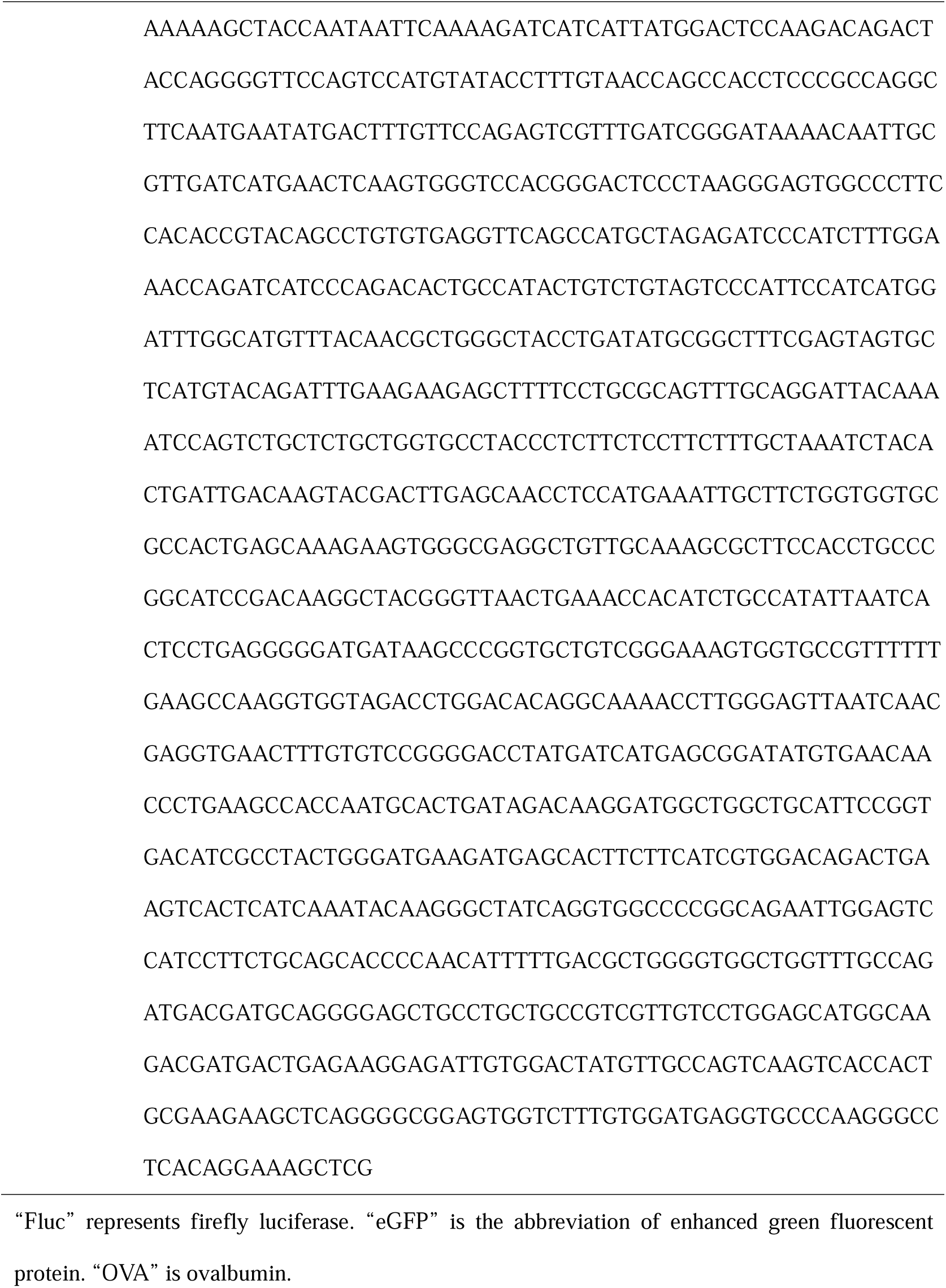
The synthetic design sequence (SDS) of in vitro transcribed mRNA was adopted in this study.

## Supplementary Figures

**Supplementary Figure 1.**
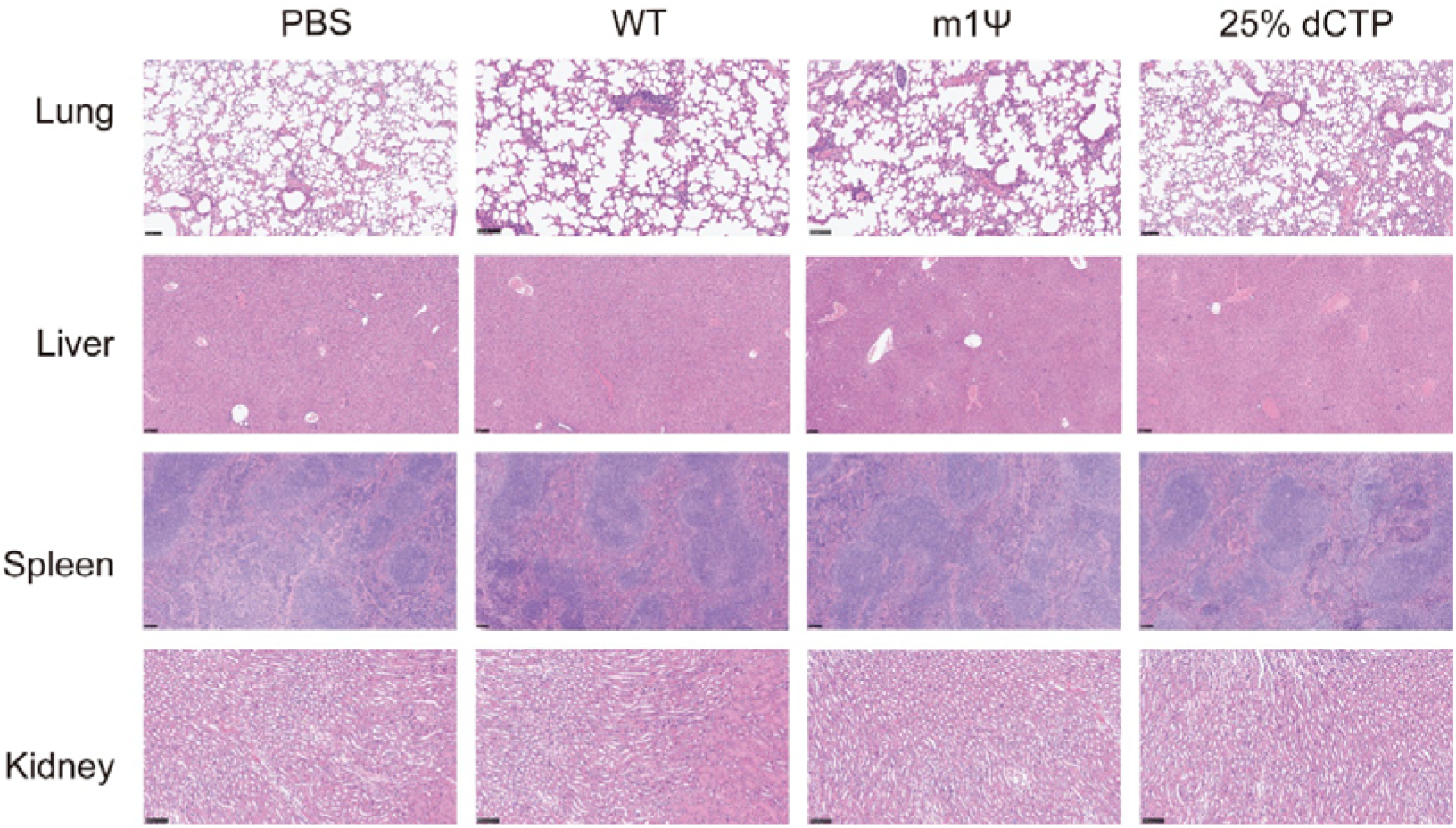
Representative H&E staining images of major organs, including lung, kidney, liver and spleen, collected after treatment with PBS, WT IVT-mRNA, m1Ψ-modified IVT-mRNA or 25% dCTP-modified IVT-mRNA. No obvious pathological abnormalities or treatment-associated tissue injury were observed in the 25% dCTP group compared with WT and m1Ψ groups. Scale bars: 50 μm.

**Supplementary Figure 2.**
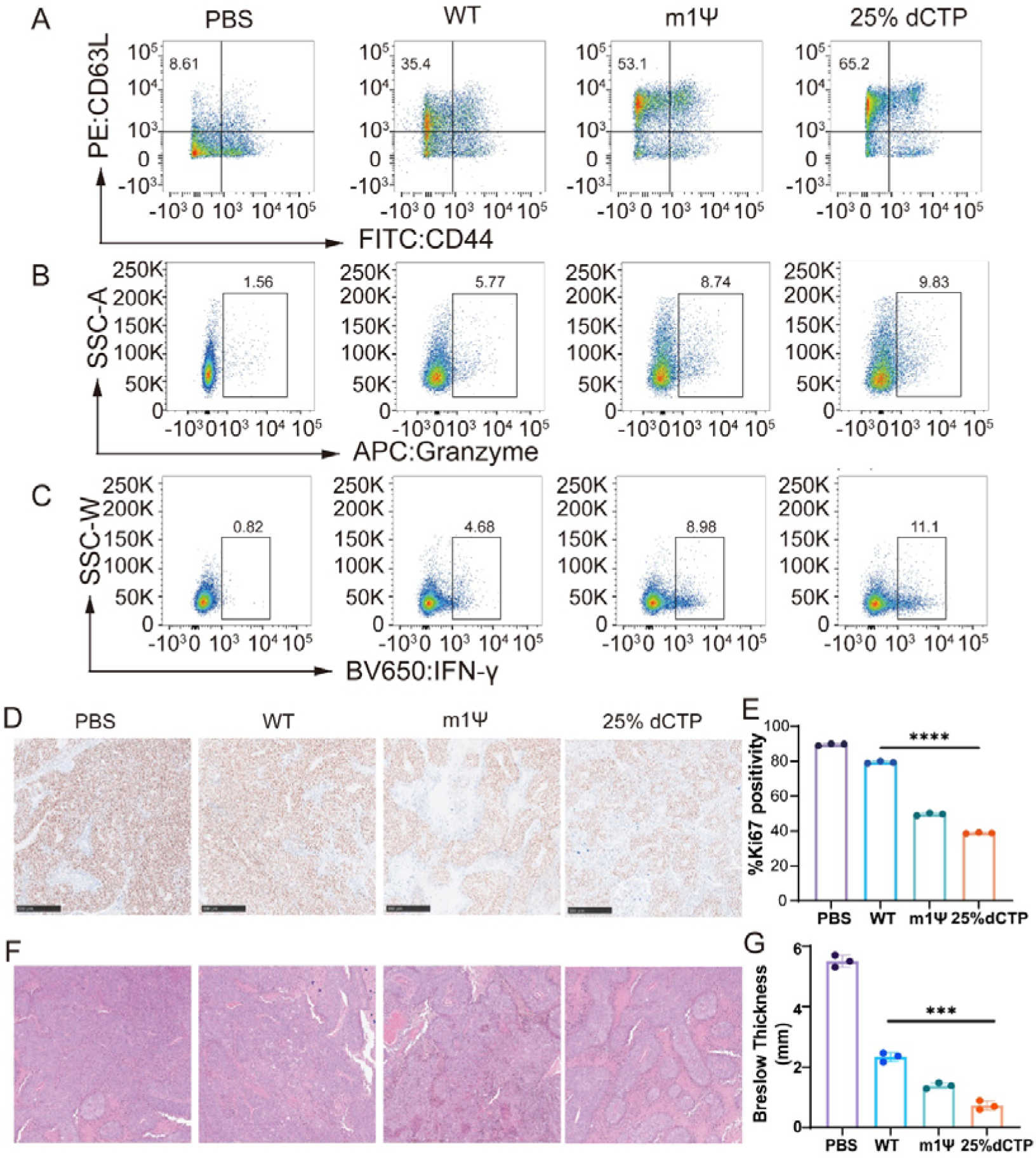
(A-C) Representative flow-cytometry plots of draining inguinal lymph nodes from B16-OVA tumor-bearing mice treated with PBS, WT IVT-mRNA, m1Ψ-modified IVT-mRNA or 25% dCTP-modified IVT-mRNA. Cells were analyzed for CD44 and CD62L expression, granzyme B production and IFN-γ production. Compared with WT and m1Ψ controls, 25% dCTP-modified IVT-mRNA induced higher frequencies of CD44□CD62L□T cells, granzyme B□cells and IFN-γ cells. (D) Representative Ki67 immunohistochemistry images of tumor sections from each treatment group. (E) Quantification of Ki67 positivity in tumor sections. (F) Representative H&E staining images of tumor sections from each treatment group. (G) Quantification of tumor thickness in H&E-stained tumor sections. Data are presented as mean ± SD.

## References

1. E. Rohner, R. Yang, K. S. Foo, A. Goedel, K. R. Chien, Unlocking the promise of mRNA therapeutics. Nature biotechnology 40, 1586–1600 (2022).

2. N. Pardi, M. J. Hogan, F. W. Porter, D. Weissman, mRNA vaccines — a new era in vaccinology. Nature Reviews Drug Discovery 17, 261–279 (2018).

3. C. Liu, et al., mRNA-based cancer therapeutics. Nature Reviews Cancer 23, 526–543 (2023).

4. J. R. Mascola, A. S. Fauci, Novel vaccine technologies for the 21st century. Nature Reviews Immunology 20, 87–88 (2020).

5. E. Callaway, M. Naddaf, Pioneers of mRNA COVID vaccines win medicine Nobel. Nature 622, 228–229.

6. K. Karikó, M. Buckstein, H. Ni, D. Weissman, Suppression of RNA Recognition by Toll-like Receptors: The Impact of Nucleoside Modification and the Evolutionary Origin of RNA. Immunity 23, 165–175 (2005).

7. K. Karikó et al., Incorporation of Pseudouridine Into mRNA Yields Superior Nonimmunogenic Vector With Increased Translational Capacity and Biological Stability. Molecular Therapy 16, 1833–1840 (2008).

8. D. Liu, H. Chen, A. Pan, X. Wang, Beyond the Sequence: Chemical and Topological Design and Innovations in mRNA Therapeutics. Chemical Reviews 126, 3907–3956 (2026).

9. X. Hou, T. Zaks, R. Langer, Y. Dong, Lipid nanoparticles for mRNA delivery. Nature Reviews Materials 6, 1078–1094 (2021).

10. L. L. Y. Ho, G. H. A. Schiess, P. Miranda, G. Weber, K. Astakhova, Pseudouridine and N1-methylpseudouridine as potent nucleotide analogues for RNA therapy and vaccine development. RSC chemical biology 5, 418–425 (2024).

11. B. Alberts, D. Bray, J. Lewis, M. Raff, K. Roberts, J. D. Watson, Molecular Biology of the Cell (Garland Publishing, ed. 2, 1989).

12. G. Hayson, Advanced Molecular Biology: Exploring the Intricacies of DNA, RNA, and Protein Functions (Springer, 2025).

13. J. Wang et al., The role of sequence context, nucleotide pool balance and stress in 2′-deoxynucleotide misincorporation in viral, bacterial and mammalian RNA. Nucleic acids research 44, 8962–8975 (2016).

14. R. W. Siegel, L. Bellon, L. Beigelman, C. C. J. J. o. v. Kao, Use of DNA, RNA, and chimeric templates by a viral RNA-dependent RNA polymerase: evolutionary implications for the transition from the RNA to the DNA world. Journal of Virology 73, 6424–6429 (1999).

15. N. Zenkin, Y. Yuzenkova, K. J. S. Severinov, Transcript-assisted transcriptional proofreading. Science 313, 518–520 (2006).

16. J. R. Wyatt, G. T. Walker, Deoxynucleotide-containing oligoribonuccleotide duplexes: stability and susceptibility to RNase V 1 and RNase H. Nucleic acids research 17, 7833–7842 (1989).

17. E. W. Frederick, U. Maitra, J. Hurwitz, The Role of Deoxyribonucleic Acid in Ribonucleic Acid Synthesis: XVI. The purification and properties of ribonucleic acid polymerase from yeast: preferential utilization of denatured deoxyribonucleic acid as template. Journal of Biological Chemistry 244, 413–424 (1969).

18. G. V. Paddock, H. C. Heindell, W. Salser, Deoxysubstitution in RNA by RNA polymerase in vitro: a new approach to nucleotide sequence determinations. Proceedings of the National Academy of Sciences of the United States of America 71, 5017–5021 (1974).

19. X. Cao, Y. Zhang, Y. Ding, Y. Wan, Identification of RNA structures and their roles in RNA functions. Nature reviews. Molecular cell biology 25, 784–801 (2024).

20. M. J. Hammerling, A. Krüger, M. C. Jewett, Strategies for in vitro engineering of the translation machinery. Nucleic acids research 48, 1068–1083 (2020).

21. D. Temiakov et al., Structural basis for substrate selection by T7 RNA polymerase. Cell 116, 381–391 (2004).

22. S. O. Gudima et al., Synthesis of mixed ribo/deoxyribopolynucleotides by mutant T7 RNA polymerase. FEBS letters 439, 302–306 (1998).

23. D. Zhang et al., Machine learning engineered PoLixNano nanoparticles overcome delivery barriers for nebulized mRNA therapeutics. bioRxiv 2024.2011.2003.621713 (2024). doi: 10.1101/2024.11.03.621713

24. L. Xu et al., Host Microenvironment Reprogramming by Saccharides Overcomes Lung Barriers for mRNA Therapeutics. bioRxiv 2025.2007.2014.664686 (2025). doi: 10.1101/2025.07.14.664686

25. U. Sahin, K. Karikó, Ö. Türeci, mRNA-based therapeutics — developing a new class of drugs. Nature Reviews Drug Discovery 13, 759–780 (2014).

26. B. Rozman et al., N1-Methylpseudouridine directly modulates translation dynamics. Nature 651, 533–541 (2026).

27. M. Bérouti et al., Pseudouridine RNA avoids immune detection through impaired endolysosomal processing and TLR engagement. Cell 188, 4880–4895.e4815 (2025).

28. S. Schiffers et al., N4-Acetylcytidine enhances synthetic mRNA translation yield and fidelity. Nature, (2026). doi: 10.1038/s41586-026-10729-8

29. J. Rudert et al., Rational design of a modular mRNA vaccine platform for rapid adaptation to SARS-CoV-2 variants. Scientific reports 16, 17178 (2026).

30. J. Zhang et al., Airway-applied mRNA vaccine needs tailored sequence design and high standard purification that removes devastating dsRNA contaminant. Molecular therapy : the journal of the American Society of Gene Therapy 33, 4193–4215 (2025).

31. Q. Xiao et al., Efficient Transfection of In vitro Transcribed mRNA in Cultured Cells Using Peptide-Poloxamine Nanoparticles. Journal of visualized experiments 17.186 (2022).

32. S. Guan et al., Self-assembled peptide-poloxamine nanoparticles enable in vitro and in vivo genome restoration for cystic fibrosis. Nature nanotechnology 14, 287–297 (2019).

